# Constitutive signaling by the C-terminal fragment of polycystin-1 is mediated by a tethered peptide agonist

**DOI:** 10.1101/2021.08.20.457171

**Authors:** Brenda S. Magenheimer, Ericka Nevarez Munoz, Jayalakshmi Ravichandran, Robin L. Maser

## Abstract

Mutation of the *PKD1* gene, encoding polycystin-1 (PC1), is the primary cause of autosomal dominant polycystic kidney disease. PC1 is an 11-transmembrane domain protein that binds and modulates the activity of multiple heterotrimeric G protein families and is thought to function as a non-canonical G protein-coupled receptor (GPCR). PC1 shares a conserved <u>G</u>PCR <u>a</u>utoproteolysis <u>in</u>ducing [GAIN] domain with the adhesion family of GPCRs, that promotes an auto-catalytic, cis-cleavage at the GPCR proteolysis site (GPS) located proximal to the first transmembrane domain. GPS cleavage divides these receptors into two associated ‘subunits’, the extracellular N-terminal (NTF) and transmembrane C-terminal (CTF) fragments. For the adhesion GPCRs, removal of the NTF leads to activation of G protein signaling as a result of the exposure and subsequent intramolecular binding of the extracellular N-terminal stalk of the CTF, i.e., the tethered cryptic ligand or tethered agonist model. Here, we test the hypothesis that PC1-mediated signaling is regulated by an adhesion GPCR-like, tethered agonist mechanism. Using cell-based reporter assays and mutagenesis of PC1 expression constructs, we show that the CTF form of PC1 requires the stalk for signaling activation and synthetic peptides derived from the PC1 stalk sequence can re-activate signaling by a ‘stalk-less’ CTF. In addition, we demonstrate that ADPKD-associated missense mutations within the PC1 stalk affect signaling and can inhibit GPS cleavage. These results provide a foundation for beginning to understand the molecular mechanism of G protein regulation by PC1 and suggest that a tethered agonist-mediated mechanism can contribute to PKD pathogenesis.

**SIGNIFICANCE STATEMENT:** Mutations of the PKD1 gene, encoding polycystin-1, are the predominant cause of autosomal dominant polycystic kidney disease (ADPKD), a systemic disease that is the 4^th^ leading cause of kidney failure. Polycystin-1 functions as an atypical GPCR capable of binding or activating heterotrimeric G proteins, which is essential for preventing renal cystogenesis. However, little is known regarding its regulation. Polycystin-1 shares structural features with the Adhesion family of GPCRs. In this work, we combined mutagenesis and cellular signaling assays which demonstrated that constitutive activation of signaling by polycystin-1 involves an Adhesion GPCR-like molecular mechanism. This study provides new knowledge regarding the structure-function relationships of polycystin-1 which will stimulate additional areas of investigation and reveal novel avenues of therapeutic intervention for ADPKD.

## INTRODUCTION

Polycystin-1 (PC1) is the product of the PKD1 gene that is mutated in 85% of the cases of autosomal dominant polycystic kidney disease (ADPKD) (1). ADPKD is a systemic disorder that affects one in every 500-1,000 individuals and is typified by bilaterally enlarged kidneys with hundreds of fluid-filled, epithelial cell-lined cysts that often lead to renal failure. PC1 is a multi-faceted protein thought to function as one subunit of a heteromeric receptor-ion channel complex formed with PC2 (2-5), the PKD2 gene product. PC1 is composed of an N-terminal, extracellular region (ECR) of >3,000 residues, 11 transmembrane (TM) domains and a short, cytosolic C-terminal tail (C-tail) (4, 6). The PC1 ECR consists of multiple interaction domains, reported to bind basement membrane components and Wnt ligands, and to mediate homotypic interactions between cells (7-10). The C-tail of PC1 has been implicated in a number of signaling cascades (11), including pathways involving heterotrimeric G proteins. Members of multiple G alpha protein families can bind to the PC1 C-tail (12-14) that harbors a conserved G-protein activation motif within its heterotrimeric G-protein binding domain (12). Both inhibition (by sequestration) and activation of Gα- and Gβγ-mediated signaling with effects on apoptosis, ion channel activity and transcription factor activation (AP-1, NFAT) have been reported for PC1 in cell-based assay systems (15-22). Recent *in vivo* studies have shown that PC1-dependent regulation of G protein signaling is necessary for preventing cyst formation in the kidneys of mice and in the developing pronephros of *Xenopus* embryos (14, 23, 24). Together, these observations support a critical function of PC1 as a novel and non-canonical GPCR important in ADPKD pathophysiology. However, the molecular mechanism by which PC1 mediates G protein regulation is currently unknown.

Intriguing similarities between PC1 and the unusual Adhesion/B2 family of GPCRs (AdGPCRs) have been noted (25, 26). One shared characteristic is the relatively large ECR composed of multiple adhesive domains (26, 27) and a conserved <u>G</u>PCR <u>p</u>roteolysis <u>s</u>ite (GPS) motif located near the first TM domain (28, 29). The GPS is embedded within a conserved <u>G</u>PCR <u>a</u>utoproteolysis <u>in</u>ducing [GAIN] domain whose unique structure is responsible for catalyzing an autoproteolytic cleavage between its final two β-strands at the GPS HL/T consensus site (30). GPS cleavage generates an extracellular N-terminal (NTF) and a membrane-embedded C-terminal (CTF) fragment, or subunits, that remain non-covalently associated due to extensive hydrogen bonding between the final (13^th^) β-strand and surrounding β-strands 6, 9 and 12. For AdGPCRs, the NTF has been shown to be involved in regulation of G protein signaling by the CTF via multiple potential mechanisms (27). In the cryptic tethered agonist (TA) model (31, 32), GPS cleavage and dissociation of the NTF subunit exposes β-strand 13, located at the beginning of the N-terminal stalk of the CTF subunit, which is envisioned to embed itself within the 7-TM helical bundle, thus leading to conformational changes and activation of heterotrimeric G proteins. Based on this scenario, β13 and/or the N-terminal stalk of the CTF are also referred to as the *Stachel* sequence (for stinger) (32). We hypothesized that PC1-mediated signaling would be regulated by a similar tethered cryptic agonist-like mechanism. Using a cell-based system with transient expression of wild type (WT) and mutant forms of PC1, we show that constitutive activation of a NFAT promoter-luciferase reporter is dependent on the stalk region of the PC1 CTF subunit, which can be substituted for *in trans* by soluble peptides derived from the stalk region. In addition, we identify ADPKD-associated missense mutations within the stalk that play essential roles in GPS cleavage and/or activation of the PC1 CTF subunit.

## RESULTS

### FL and CTF expression constructs of PC1 constitutively activate an NFAT promoter-luciferase reporter to different extents

Previous studies showed activation of AP-1 and NFAT promoter-luciferase reporters by a membrane-directed expression construct encoding the C-terminal 222 aa-residues of PC1 (C-tail) via Gαi-Gβγ*/*JNK and Gαq/PLC/Ca2+ signaling pathways, respectively (15, 17). We first determined if full-length [FL] PC1 is capable of signaling to the AP-1 or NFAT reporters by transient transfection of HEK293T cells with FL-PC1 expression constructs encoding either human (h) or mouse (m) PC1 (**Fig. 1A**). Transfection of different amounts of mFL-PC1 or hFL-PC1 (data not shown) construct led to a ‘dose’-dependent increase in NFAT reporter activation in comparison to the empty expression vector (ev) control (**Fig. S1A**). Western blot of transfected cell lysates demonstrated the presence of NTF, CTF and uncleaved FL (FL^U^) forms of FL-PC1 (**Fig. S1B-D**). Increases in NFAT reporter activation appeared to correspond with the increases in CTF expression levels. Neither mFL-PC1 nor hFL-PC1 consistently activated the AP-1 reporter (data not shown), therefore we continued using only the NFAT reporter in further analyses.

**Figure 1.**
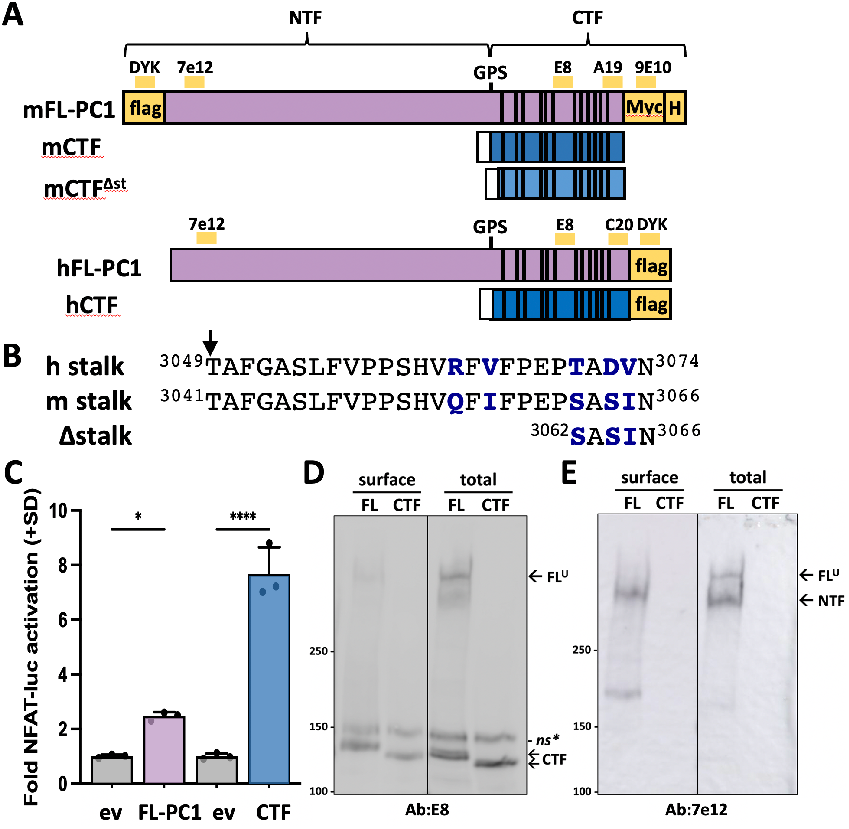
CTF has greater signaling activity than FL-PC1 to NFAT reporter. (**A**) Illustration of the full-length [FL] and C-terminal fragment (CTF) expression constructs of PC1 from mouse (m) and human (h). GPS, autocatalytic cleavage site that generates NTF and CTF ‘subunits’ from FL-PC1. Yellow bars, epitope location for antibodies (Ab) used in this study; white rectangle, CD5 signal sequence; vertical lines, 11 TM domains; H, 6xHis. (**B**) Alignment of stalk region sequences of human and mouse PC1. mCTF^Δst^ has a 21-residue deletion from the N-terminal end of the stalk region. Arrow, GPS cleavage site. Non-identical residues shown in bolded blue. (**C**) Cells were transfected with mFL-PC1, mCTF, or corresponding equimolar amounts of empty expression vector (ev) control DNA. Graph shows the mean fold NFAT-luc activation (+SD) for each condition (n=3 wells/condition). Result is representative of 3 independent experiments. *, p<0.05; ****, p<0.0001. (**D**,**E**) Western blot of total cellular and biotinylated surface PC1 proteins from transfection in (**C**) probed with E8 (D) or 7e12 (E). Migration of CTF band differs between mFL-PC1 and mCTF due to the C-terminal Myc and His tags of the mFL-PC1 construct. *, non-specific band. Specificity of E8 Ab demonstrated in Fig. S1B. Line between lanes of a blot indicates site of removal of non-relevant intervening lanes.

AdGPCRs that utilize a TA mechanism exhibit greater signaling activity when NTF sequences are deleted from the expression construct, i.e., ΔNTF or CTF alone (32, 34-37). To determine if PC1-mediated activation of the NFAT reporter exhibits this characteristic, CTF-only expression constructs of human (h) and mouse (m) PC1 were made consisting of a CD5 signal peptide sequence (**Fig. 1A**) preceding the stalk region sequences of hCTF and mCTF (i.e., initiating with residue T3049 or T3041, respectively) (**Fig. 1B**). Transfection conditions were established to achieve similar levels of expression from the FL and CTF constructs in the same transfection experiment in order to directly compare their abilities to activate the NFAT reporter. Both FL and CTF expression constructs of mPC1 increased NFAT reporter activity multiple-fold over the ev control, albeit to different extents (**Fig. 1C**). Similar results were obtained using hFL-PC1 and hCTF expression constructs (**Fig. S2A**). Western blot analysis of total cell lysates with antibodies recognizing epitopes within the NTF or the CTF showed that equivalent amounts of the CTF form were expressed in the FL-PC1- and CTF-transfected cells (**Fig. 1D**, total lanes) and that FL-PC1 underwent efficient GPS cleavage, i.e., level of NTF was much greater than FL^U^ (**Fig. 1E**, total lanes), respectively. Surface biotinylation experiments performed in parallel showed that similar levels of the CTF subunit reached the cell surface from transfected FL-PC1 and CTF constructs (**Fig. 1D**, surface lanes). As expected, the NTF and CTF subunits were much more abundant than FL^U^ at the plasma membrane (**Fig. 1E**), demonstrating the specificity of the procedure. Similar results were obtained with surface biotinylation analyses of cells transfected with the hFL-PC1 construct (**Fig. S2C**; note the additional controls included). Altogether, these results demonstrate that when absent of the NTF subunit, the PC1 CTF has greater constitutive activity than cleaved FL-PC1 and is able to traffic to the cell surface.

### The stalk is required for constitutive NFAT reporter activation by PC1 CTF

To determine if the stalk is acting as an agonist in the CTF construct, a ‘stalk-less’ version lacking the first 21 residues was generated, designated CTF^Δst^ (**Fig. 1A,B**), and its signaling ability was directly compared to CTF. The CTF^Δst^ mutant exhibited a dramatic loss of NFAT reporter activation that was essentially reduced to ev control levels (**Fig. 2A**). Both steady state and cell surface expression levels of CTF^Δst^ were comparable to CTF (**Fig. 2 B-E**), which suggests that neither protein instability nor loss of membrane trafficking was responsible for the inability of CTF^Δst^ to activate the NFAT reporter. These results are consistent with the PC1 CTF stalk acting as a tethered peptide agonist.

**Figure 2.**
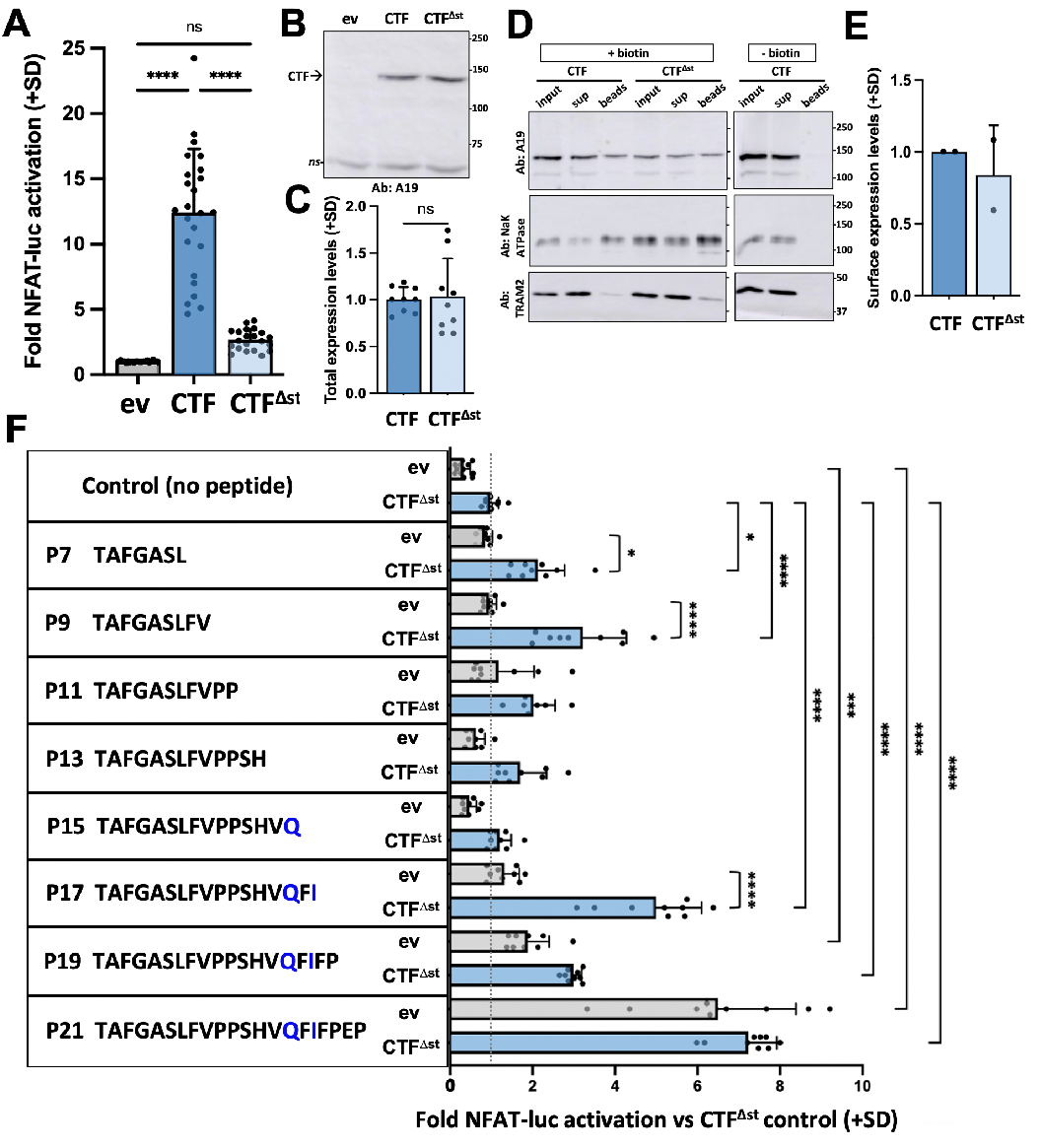
The stalk region is necessary for signaling by PC1 CTF. (**A**) Activation of the NFAT-luc reporter by transfected mCTF or mCTF^Δst^ expression constructs relative to empty vector (eV). Shown are the means + standard deviation [SD] of 3 wells/construct from each of 7 independent experiments. (**B**) Representative Western blot of total cell lysates from one of the experiments in (A), probed with A19. ns, non-specific. (**C**) Summary of the total expression levels (means +SD) of CTF^Δstk^ relative to CTF from the experiments in (A). (**D**) Representative Western blot of a surface biotinylation experiment for mCTF and mCTF^Δstk^. Experiment included a ‘-biotin’ control with CTF-transfected cells for which the NHS-biotin reagent was omitted from the procedure. Aliquots of the total cell lysate (input; after biotinylation and before neutravidin pulldown), supernatant (sup; following neutravidin bead removal), and biotinylated cell surface proteins bound to neutravidin beads (beads) representing 10%, 10% and 70% of each sample, respectively, were analyzed. Top blots were probed with A19 and then stripped and reprobed for NaKATPase or the ER resident protein, TRAM2. (**E**) Summary of cell surface expression levels of CTF^Δstk^ relative to CTF (means +SD) from two separate experiments. (**F**) Stalk peptide treatment of ev- or m CTF^Δstk^-transfected cells. Sequences of stalk-derived peptides P7-P21 are shown. Graph represents the fold NFAT-luc activation for both eV- (gray bars) and CTF^Δst^- (blue bars) transfected cells relative to the CTF^Δst^ control after 24 hr treatment with or without peptide. Results are the means (+SD) of 3 separate experiments, each with 3 wells/condition. *, p < 0.05; ***, p = 0.0001; ****, p < 0.0001. Analysis by 1-way ANOVA with Tukey-Kramer post-test.

### Synthetic, stalk-derived peptides re-activate NFAT reporter by CTF^Δst^ *in trans*

To further verify the agonist property of the PC1 CTF stalk, peptides (P) consisting of the N-terminal 7, 9, 11, 13, 15, 17, 19 or 21 residues of the stalk sequence from mPC1 followed by a short (7-residue) hydrophilic sequence were synthesized and used to treat ev- or mCTF^Δst^-transfected cells. The ‘no peptide’ control consisted of addition of culture medium only. Treatment with P7, P9 or P17 resulted in significant activation of the NFAT reporter in CTF^Δst^-transfected cells in comparison to their corresponding peptide-treated, ev-transfected controls and to the CTF^Δst^ with no peptide treatment control (**Fig. 2F**). Treatment with either P19 or P21 significantly increased NFAT reporter activity in both ev- and CTF^Δst^-transfected cells in comparision to the no peptide treatment controls, which showed that P19- and P21-mediated reporter activation was not dependent on the presence of CTF^Δst^, thereby suggesting that an endogenous protein(s) was activated by these longer peptides. Treatment with P19 or P21 also resulted in ‘rounding up’ of both ev- and CTF^Δst^-transfected cells, which was not observed with the other stalk peptides (data not shown). These results demonstrate that stalk-derived peptides P7, P9 and P17 specifically stimulated signaling by the stalkless CTF^Δst^ mutant protein to the NFAT reporter, which is consistent with the PC1 CTF stalk region harboring agonist activity.

### Missense variants within the stalk region alter NFAT activation by PC1 CTF

The PKD Mutation Database (pkdb.mayo.edu) lists a total of 11 missense variants located at 7 positions within the PC1 stalk region (**Fig. 5A**), one of which, F3066L, is considered a common benign polymorphism (38, 39). To determine if the ADPKD-associated mutations affect agonist activity of the stalk, we generated mutant expression constructs of hCTF with a single variant from each of the other 6 sites (**Fig. 3A**) and determined their effect on signaling. NFAT reporter activation was significantly reduced for CTF mutants G3052R, V3057M, R3063C and N3074K, significantly increased by P3069L, and not affected by the E3068D mutant in comparison to WT CTF (**Fig. 3B**). Western blot analyses showed that none of the missense mutations had a significant effect on either steady state or cell surface expression levels in comparison to WT CTF (**Fig. 3C-F**). We also generated mutant expression constructs for mCTF with G3044R and Q3055C variants, synonymous to hG3052R and hR3063C mutations, and determined their signaling ability. Each mutation significantly reduced NFAT reporter activation (**Fig. S3**). These results suggest that TA activity of the PC1 CTF is dependent on specific residues and/or positions within the stalk region.

**Figure 3.**
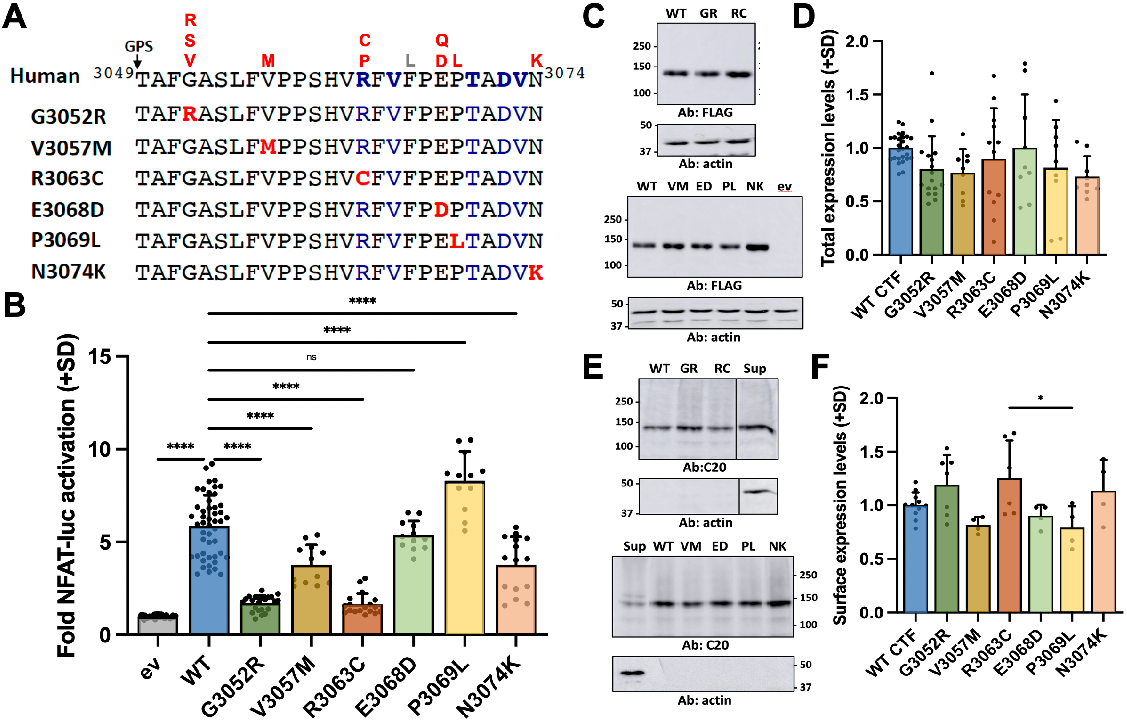
ADPKD-associated missense mutations within the stalk region affect signaling by CTF. (**A**) Top, locations of ADPKD missense amino acid mutations (red) in the PC1 stalk region reported in PKD Mutation Database (pkdb.mayo.edu). F3066L (in gray) is a frequently reported benign polymorphism. Below, stalk mutant CTF expression constructs generated for analysis. Mutated residues (red); residues differing from mouse PC1 stalk (blue). (**B**) Activation of the NFAT reporter by empty vector (ev), wild type (WT) and mutant CTF expression constructs. Shown are the means (+ standard deviation, SD) of 4-12 independent transfection experiments each with 3 technical replicates per construct. (**C**) Western blot analyses of total cell lysates from ev-, WT-, and ADPKD mutant CTF-transfected cells. Blots were probed with anti-FLAG antibody to detect hCTF and re-probed with antibody against actin. Mutants are indicated by abbreviations: GR, G3052R; RC, R3063C; VM, V3057M; ED, E3068D; PL, P3069L; and NK, N3074K. Blot is representative of the transfections in (B). (**D**) Graph summarizes the means (+SD) for total cell expression levels of ADPKD stalk mutants relative to WT CTF from the transfection experiments in (B). (**E**) Representative Western blots of the streptavidin-pull downs of surface biotinylated proteins probed with C20 antibody (Ab) to detect hCTF and then re-probed for actin. Sup, aliquot of the supernatant from the streptavidin pull-down of WT CTF. The line between lanes of a blot indicates removal of non-relevant intervening lanes. (**F**) Graph shows the surface expression levels (means +SD) from 2-4 separate transfection experiments. *, P < 0.02; **, P < 0.002; ***, P < 0.0001.

### An ADPKD missense mutation in the stalk region alters GPS cleavage of full-length PC1

GPS cleavage is thought to be essential for the full functionality of PC1 since mutations that interfere with cleavage result in PKD in mice (40, 41). To determine if the selected ADPKD mutations affect GPS cleavage, hFL-PC1 expression constructs with each of the missense mutations were generated (**Fig. 4A**) and transfected into 293T cells. For comparison, expression constructs for WT FL-PC1 and the known GPS cleavage mutant, FL-PC1-T3049V (28), were also included. As expected, Western blot analyses showed efficient GPS cleavage of WT FL-PC1 and a complete lack of cleavage for FL-PC1-T3049V, and in addition, for the G3052R mutant (**Fig. 4B,C**). For each of the other ADPKD stalk mutants, both NTF and CTF subunits were detected at levels similar to those of WT FL-PC1. Together with the effects on NFAT reporter activation by stalk mutant CTF expression constructs (**Fig. 3B**), these results suggest that the G3052R substitution may be pathogenic due to its inhibition of PC1 GPS cleavage, while the pathogenicity of ADPKD-associated mutations V3057M, R3063C, P3069L and N3074K may result from their effects on the activity and/or structure of the TA of the CTF subunit.

**Figure 4.**
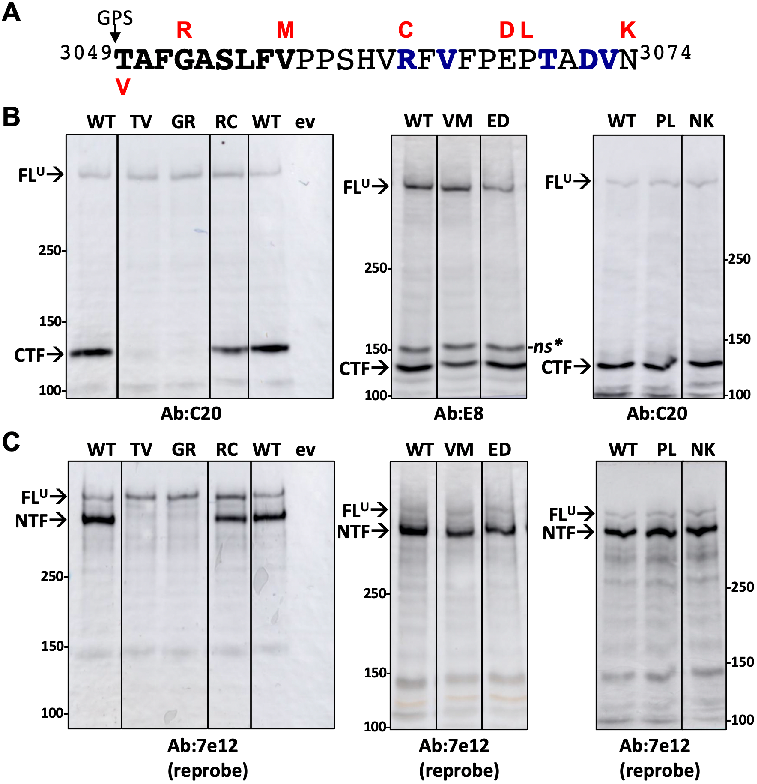
Effect of ADPKD-associated stalk mutations on GPS cleavage. (**A**) Human PC1 stalk region sequence with the ADPKD missense mutations tested in FL-PC1 shown above the stalk sequence in red. Residues in blue indicate those that differ in mouse PC1 stalk. Val substitution at T3049 is known to inhibit GPS cleavage. (**B**,**C**) Western blot analyses of human PC1 stalk region mutants following expression in 293T cells. (**B**) Blots were probed with antibodies recognizing epitopes within the CTF. (**C**) Same blots from (B) were stripped and reprobed with antibody 7e12. Non-cleaved, full-length [FL] and NTF and CTF cleavage subunits of PC1 are indicated. The line between lanes of a blot indicates site of removal of non-relevant intervening lanes.

## DISCUSSION

This work represents the first demonstration that PC1, a non-canonical, GPS-cleaved, 11-TM GPCR, utilizes an AdGPCR-like, tethered peptide agonist for constitutive signaling. We used transient transfection of a 4xNFAT response element-luciferase reporter previously shown to be activated by the PC1 C-tail via the canonical Gq-PLC-Ca^2+^ signaling pathway (17) for these studies. Here, we show that both FL-PC1 and the CTF subunit alone (ΔNTF) constitutively activate the NFAT reporter when ectopically expressed in HEK293T cells. Direct comparisons of CTF and FL-PC1 demonstrated a ≥ 3-fold activation of the NFAT reporter by the CTF subunit of both human and mouse PC1. CTF-mediated activation of the reporter was dependent on its N-terminal, extracellular stalk, as shown by the loss of reporter activity with a stalk-deletion expression construct (CTF^Δst^) and by the ability of synthetic, stalk-derived peptides to reactivate signaling by CTF^Δst^ *in trans*. Furthermore, the potential pathogenic relevance of a TA-mediated activation mechanism for PC1 was supported by the demonstration that certain ADPKD-associated missense mutations within the stalk region altered the signaling capability of the CTF subunit and/or inhibited GPS cleavage of FL-PC1.

The presence of the GAIN domain and GPS cleavage in both AdGPCRs and PC1 (30), coupled with their abilities to activate signaling via heterotrimeric G proteins, has led to proposals of a shared functional mechanism for these two unusual protein families (25, 26, 42). Up until now, however, direct experimental evidence demonstrating an AdGPCR-like mechanism for PC1 has been lacking, although a few previous observations for PC1 are consistent with such an activation mechanism and are reminiscent of results obtained for AdGPCRs (18, 22, 43, 44). One of these studies used a CTF-like expression construct of PC1, called PC1-11TM, and demonstrated its ability to activate the NFAT reporter (17). However, PC1-11TM was a chimeric protein that included 32 residues preceding the GPS cleavage site fused to the CH_2_-CH_3_ domains of human IgG as its ectodomain, and it did not undergo cleavage, most likely due to its lacking additional portions of the GAIN domain required for self-catalyzed proteolysis (30). Likewise, PC1-11TM’s incomplete GAIN domain structure would not be expected to completely ‘encrypt’ the stalk, and thus may have allowed TA-mediated signaling. Therefore, to definitively test the existence of a TA mechanism for PC1, we generated ‘authentic’ versions of the PC1 CTF beginning with residue hT3049/mT3041 and cloned in frame with the cleavable CD5 signal peptide sequence (6, 45) (**Fig. 1A,B**). As such, this work provides the first direct investigation of a TA-like mechanism for PC1-mediated signaling.

Our experimental results strongly support the conclusion that the constitutive signaling activity of PC1 is due to a cryptic, TA within the N-terminal stalk region of its CTF subunit. First of all, direct comparison of the signaling activities of CTF and FL-PC1 showed that the CTF construct consistently led to a greater fold activation of the NFAT reporter under conditions of similar levels of total and cell surface expression (**Fig. 1**). Similar observations have been reported for a number of Ad-GPCRs (27), which were originally interpreted as an ability of the associated NTF subunit to inhibit CTF-G protein signaling via interactions that stabilized an inactive conformation of the CTF (46). Second, deletion of the N-terminal, 21 residues of the stalk in the CTF^Δstk^ mutant resulted in the loss of NFAT reporter activation, although CTF^Δstk^ was expressed and reached the cell surface at levels comparable to those of WT CTF (**Fig. 3**). Third, soluble peptides consisting of N-terminal sequences of the PC1 stalk were capable of bestowing NFAT reporter activation specifically to CTF^Δstk^-transfected cells. All three of these features have been demonstrated for many AdGPCRs and are considered requirements for a cryptic TA (31, 32). Once the CTF stalk was discovered as a tethered agonist for the AdGPCRs, coupled with knowledge of the GAIN domain structure (30), NTF-mediated inhibition of signaling was attributed to its sequestering of the N-terminal portion of the CTF stalk, the β13 strand or the *Stachel* sequence. This sequestering occurs within the interior of the GAIN domain due to extensive hydrogen-bonding interactions, which also maintain the non-covalent association between NTF and CTF subunits of GPS-cleaved AdGPCRs (30). Likewise, the differential signaling ability of the PC1 CTF and FL-PC1 reported in this study could be a consequence of the non-covalent association of NTF and CTF subunits following GPS cleavage, as has been previously reported (28, 47). The cryptic TA model implies that the PC1 NTF subunit would have to dissociate in order for the stalk to activate signaling by the CTF to the NFAT reporter. It is conceivable that removal of the PC1 NTF could occur *in vivo* by mechanical means, or through its binding to specific ECM constituents or to itself via the PKD repeats in *cis* or *trans* conformations (8, 48-50). Similar scenarios have been demonstrated for various Ad-GPCRs (33, 51-54). In support of the ability for PC1 NTF-CTF subunits to dissociate, it has been shown that treatment of cells with an alkaline solution will lead to release of the NTF from the CTF in the membrane (55), and there is evidence that the PC1 CTF subunit exists by itself, i.e., dissociated from NTF, *in vivo* (47). Interestingly though, flexibility within the GAIN domain was recently shown to allow transient exposure of parts of the encrypted TA of a number of AdGPCRs (56), which may underlie the proposed allosteric activation mechanism whereby ligand-binding to the ectodomain enables signaling in absence of NTF removal and may account for the signaling ability of some GPS cleavage-deficient AdGPCRs (57-60). It will be important to determine whether the PC1 GAIN domain also exhibits such characteristics in future investigations.

A frequency plot for the stalk region sequence of PC1 orthologues (**Fig. 5A**) reveals that the N-terminal portion is highly conserved, consisting predominantly of nonpolar residues, which bears some resemblance to the 9-residue consensus motif, or core peptide region identified at the N-terminus of AdGPCR stalks that is typically composed of β-stand 13 (**Fig. 5B**) (31, 59, 61). The relatively hydrophobic/nonpolar nature of the AdGPCR stalk core region is thought to be important for favoring its insertion into the 7-TM bundle after its exposure upon release of the NTF, i.e., the *Stachel* (or stinger) sequence (32). The 25-residue length of the PC1 stalk is also consistent with the Ad-GPCR stalks which range between ∼15-25 residues in length, in addition to its predicted ability to form secondary structure elements separated by a turn residue when separate from the GAIN domain (**Fig. 5C**) (32). Agonistic peptide sequences identified for AdGPCRs range from 7 – 19 residues in length and were derived from the N-terminus of the stalk (27). Similarly for PC1, we find that synthetic peptides consisting of the first 7-, 9- or 17-residues of the CTF stalk were agonistic (**Fig. 2**). The observation that two different lengths of stalk-derived peptides could restore NFAT reporter activation by PC1 CTF^Δst^ is somewhat unusual but might reflect the need for more than just the N-terminal *Stachel*-like sequence for ‘full’ agonist activity of the PC1 stalk. Such an idea has been proposed previously for Ad-GPCRs (32, 59, 61-63), and the presence of ADPKD-associated missense mutations located throughout the PC1 stalk region lends support to this possibility. Alternatively, in addition to the differences between the N-terminal stalk ‘consensus motifs’ of AdGPCRs and PC1 (**Fig. 5B**), our peptide data may hint at a modified mode of interaction and/or activation for this unusual 11-TM GPCR. Future work will determine the biochemical and biophysical properties of residues important for the agonist activity of the PC1 stalk.

**Figure 5.**
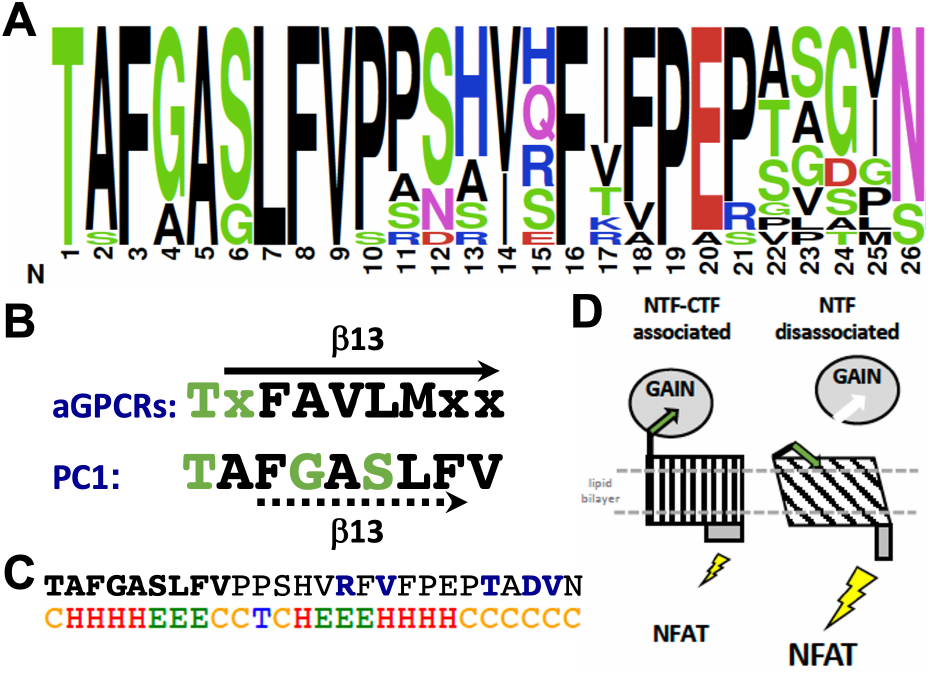
Features of the PC1 stalk region sequence and model of Adhesion GPCR-like, tethered agonist activation mechanism of PC1. (**A**) Frequency plot of the amino acid residues found at each position of the stalk and initial residue of TM 1 of PC1. Height of each residue corresponds to its frequency at that position. Color indicates chemical property of the residue: green, polar; blue, basic; red, acidic; purple, neutral; and black, hydrophobic. Plot generated by WebLogo 3 (weblogo.berkley.edu) from a multiple sequence alignment (UniProt) of 15 vertebrate species: human → pufferfish (data not shown). GPS cleavage (arrow). Identity and location of ADPKD-associated mutations are indicated above the frequency plot. (**B**) Stachel sequence consensus motif/core TA region of Adhesion GPCRs. Residues composing beta strand 13 (β13) of rat CL1/Latrophilin1/ADGRL1 are overlined with arrow, while PC1 stalk residues predicted to form β 13 (30) are underlined with a dashed arrow. (**C**) Chou-Fasman 2° structure prediction of the (exposed) hPC1 CTF stalk. C, coil; H, helix; E, sheet; T, turn. (**D**) Model of G protein signaling by PC1 as activated via a tethered (cryptic) agonist mechanism. FL-PC1 consisting of associated NTF and CTF subunits signaling has low activity. Dissociation of the NTF or its absence, as represented by the CTF expression constructs used in these studies, results in higher signaling activation due to interaction of the previously encrypted stalk/*Stachel* sequence with the remainder of the CTF.

In addition to supporting a TA mechanism for PC1-mediated signaling, our analyses of ADPKD-associated missense mutations within the PC1 stalk provide the first functional data relating to their pathogenicity that heretofore had been predicted by algorithms. For example, R3063C and P3069L, predicted as likely neutral and indeterminant, respectively (pkdb.mayo.edu), were shown to alter NFAT reporter activation by the PC1 CTF. Interestingly, P3069L led to a greater activation of the reporter than WT CTF. In this regard, it may be relevant to note that in addition to knock out, knock in missense and splicing mutations, the over-expression of *Pkd1* results in PKD in transgenic mice (64). For G3052R, which is predicted to be ‘likely pathogenic’ in the PKD Mutation Database, our analyses showed inhibition of both GPS cleavage for FL-PC1 and TA-mediated activation of CTF, thus supporting G3052R as highly likely to be pathogenic. Effects on both processes is likely a result of G3052 being located in (the predicted) β13 strand of PC1 (30), central to the GAIN domain structure, and also comprising part of the agonistic *Stachel*-like sequence at the N-terminus of the CTF stalk. In light of the demonstrated lack of effect on GPS cleavage for FL-PC1, the alteration of NFAT reporter activation by mutations V3057M, R3063C, P3069L and N3074K in the CTF expression construct may reveal a new pathogenic mechanism in ADPKD resulting from missense mutations located within the TA/stalk region.

Altogether, our data demonstrate that activation of PC1 signaling can occur via an AdGPCR-like tethered peptide agonist mechanism, consistent with the structural and functional features shared between these two protein families. We therefore propose a model for constitutive signaling by PC1 similar to the ‘cryptic tethered agonist’ model for AdGPCRs (32) in which shielding of the stalk TA by the associated NTF subunit in FL-PC1 results in a ‘weak’ signaling state, while dissociation of the PC1 NTF permits interaction of the TA stalk with the 11-TM bundle, resulting in structural conformation changes and increased signaling (**Fig. 5D**). Finally, it should be mentioned that our studies have assayed activation of a single signaling pathway reporter and it is possible that different mechanisms may be utilized by PC1 in the regulation of various G proteins and/or in activation of signaling via other pathways. Regardless, this study adds to our understanding of the structure-function relationships of PC1, provides new knowledge regarding the pathogenetic mechanisms of ADPKD and paves the way for future studies to provide further insights that could lead to the targeting of this mechanism as a potential therapy for ADPKD.

## EXPERIMENTAL PROCEDURES

### Antibodies and peptide synthesis

Antibodies used for detection of PC1 include E8-8C3C10 Ab (Baltimore PKD Center; Kerafast/EMD303), 7e12/sc-130554, C20/sc-10372 and 9E10/sc-40 (Santa Cruz Biotechnology), anti-DYKDDDDK/FLAG (Rockland; 600-401-383), and A19 (65). Additional primary antibodies included those recognizing TRAM2 (Epitomics, 3685-1), GRP78/BiP (Sigma, G9043), NaKATPase (Santa Cruz Biotechnology; sc-48345), and actin (Sigma, A2066). Secondary antibodies conjugated to HRP were purchased from Sigma or Jackson ImmunoResearch. Stalk-derived peptides were synthesized by GenScript using the Fmoc method and verified by HPLC-MS analysis.

### DNA expression constructs and cloning

Expression constructs of FL-PC1 of human (AF20) and mouse (pcDNA4/TO 3xFlag-m-PKD1-2xMyc/His) were provided by F. Qian (Addgene #21359) (28) and T. Wiembs (Addgene #83452) (66), respectively. The 3xFlag-m-PKD1-2xMyc/His insert was subcloned into pCI vector (Promega). To produce CTF expression constructs of human (h) and mouse (m) PC1, sequences starting at T3049 (human) or T3041 (mouse) and proceeding past the first TM domain were amplified by PCR from AF20 and PC1-11TM (17), respectively, using 5’-hCTFXho-BsmBI-For and 3’-BsrGIstalk-Rev primers to produce hCleavStk, and using 5’-mCleavStalkBsm-For and 3’-TMI-EcoRV primers to produce mCleavStk, each of which was joined via the BsmBI site to a PCR product encoding the CD5 signal peptide sequence (6) in pBlueScript (pBS) to generate pBS-hCD5-cleaved stalk or pBS-mCD5-cleaved stalk. The EcoRI-KpnI fragment containing hCD5-cleaved stalk was then used to replace the 5’-9.5 kb EcoRI-KpnI fragment of AF20 in pCI to generate the final pCI-hCTF expression construct, while the EcoRI-EagI fragment containing mCD5-cleaved stalk was joined to a 3.2 kb EagI-NotI fragment from PC1-11TM encoding TM2 through the C-tail to produce the final pCI-mCTF expression construct. The stalkless CTF mutant expression construct, pCI-mCTF^Δst^, starting with S3062 of the mouse PC1 stalk was generated by the same scheme except for using the 5’-mΔStalkBsmFor primer for the initial PCR. Either pBS-hCD5-cleaved stalk, pBS-mCD5-cleaved stalk, or a 516 bp BamHI-KpnI fragment from AF20 subcloned into pBS served as templates for subsequent mutagenesis of the stalk region with the QuikChangeII Site-Directed Mutagenesis Kit (Amersham). The mutagenized BamHI-KpnI fragment was used to replace the BamHI-KpnI fragment of AF20 to generate the FL-PC1 stalk mutants. PCR and mutagenesis primers were synthesized by IDT and sequences are listed in Table S1. PCR and mutagenesis products and their final constructs were confirmed by Sanger sequencing (GeneWiz). In the conduct of research utilizing recombinant DNA, the investigator adhered to NIH Guidelines for research involving recombinant DNA molecules.

### Cell culture and transient transfection

HEK293T cells (ATCC) were maintained and transiently transfected as described previously (67). Cells were passaged into 6-well plates (6 × 10^5^ cells/well; 3 wells/transfection condition) and transfected with a DNA mixture containing either the 4xNFAT or the 7xAP-1 promoter-Firefly luciferase reporter (100 ng; Stratagene), along with Renilla luciferase (50 ng of pGL4.70[*hRluc*] or 1 ng of pRL-null; Promega), and pCI expression vector encoding either FL-PC1, CTF (75 ng for signaling; 600 ng for surface biotinylation) or an equimolar amount of empty pCI vector as control. pBlueScript (Stratagene) was used to bring the total DNA amount to 8 ug. After 2.5 hrs, the DNA mixture was replaced with serum-free culture medium, and after 20-24 hrs, cells were lysed in 1X Passive Lysis buffer (PLB; Promega) supplemented with protease and phosphatase inhibitors. Firefly (Fluc)- and Renilla (Rluc)-luciferase activity in each cell lysate was determined using the Dual Luciferase Assay Kit (Promega) and a Berthold tube luminometer. NFAT-Fluc luminescence was normalized to Rluc for each well within a transfection condition, and then averaged for each condition (n =3 wells/condition). Means of normalized NFAT-Fluc with standard deviation [SD] were graphed. Signaling-transfection experiments were performed a minimum of 3 separate times each with 3 replicates/condition except where noted.

### Stalk peptide treatment

Cells were plated into 24-well plates (1.5 × 10^5^ cells/well) and transfected with CTF^Δst^ or empty pCI expression vectors, along with NFAT-Fluc and Rluc plasmids. Two hours following medium exchange, one-half of the culture medium volume was replaced with an equal volume of either serum-free medium (no peptide control) or stalk-derived peptide (2 mM in serum-free medium) and incubated overnight. In some experiments, an additional 100 ul of peptide (1 mM) was added the following morning. Cells were lysed at 24 hrs following the initial peptide or control medium addition.

### Cell surface biotinylation analyses

Surface labeling (68) was performed on intact cells 22-24 hrs post-transfection using 1.5 mg/ml PBS of the membrane-impermeable, cleavable biotin cross-linking reagent (Sulfo-NHS-SS-Biotin; Pierce) for 30 min on ice. Crosslinking was inactivated by addition of 50 mM Tris, pH 8.0. Cells were washed and then lysed in 1X PLB with protease inhibitors. A 10% aliquot of the total cell lysate was removed and saved as the input sample. NeutraAvidin-agarose beads (Pierce) were added to remove biotinylated surface proteins. The supernatant was removed and saved as the unbound cytosolic fraction (sup). A representative amount of each fraction, i.e., the total lysate (input), cytosolic (sup), and biotinylated surface proteins (beads) was analyzed by SDS-PAGE/Western blot. Cell fraction markers included NaK-ATPase (plasma membrane/surface protein), TRAM2 or BiP (ER resident proteins) and actin (cytosolic protein).

### Western blot analyses

Gel loading volumes of signaling cell lysates were calculated based on normalization to relative Rluc activity within a condition. Lysate samples were electrophoresed through 7.5% or 4.2% denaturing polyacrylamide gels and transferred to nitrocellulose membrane using the TransBlot Turbo and ReadyBlot transfer buffer (BioRad). Blots were incubated with primary antibody at 1:1,000 dilution (1:5,000 for anti-BiP; 1:10,000 for anti-TRAM2) in Tris-buffered saline (TBS; 10 mM Tris pH 7.4, 150 mM NaCl) with 0.1% Tween-20 (TBST) and 5% powdered dry milk for 14-16 hrs at 4°C. Secondary antibodies conjugated to HRP were incubated at 1:5,000 dilution in TBST/5% milk for 1 hr at room temperature. Blots were developed using a chemiluminescent substrate (Clarity; BioRad), and multiple exposures were captured for each blot with the Amersham Imager 600 with the band saturation detection mode enabled. Volume density (minus background) of immunoreactive bands was determined using ImageQuant TL (GE Healthcare). In most cases, duplicate blots were prepared and quantified to obtain an average band density relative to wild type CTF.

### Statistical analyses

Statistical analysis was performed using GraphPad Prism 9 (GraphPad Software, San Diego, CA, USA). Data are presented as means + standard deviation [SD] for bar graphs. Multiple comparisons were performed via one-way ANOVA and Tukey’s post-hoc analysis. P ≤ 0.05 was considered statistically significant.

## ACKNOWLEDGEMENTS

This work was supported by funding from the National Institutes of Health (R01 DK123590), the Department of Defense CDMRP PRMRP Discovery Award (PR160710/W81XWH-17-1-0301) and pilot grants from the School of Health Professions and the Lied Foundation at University of Kansas Medical Center to R.L.M. We sincerely thank the students and lab volunteers, including Terry Petersen, Gregory Scholtz, Arjun Ishwar, and Liz Schaffer, who contributed in various ways to the development of this work. We acknowledge the provision and/or access of PC1 expression constructs by our colleagues Feng Qian and Thomas Weimbs via Addgene.

