## Supplemental Materials for "Constitutive signaling by the C-terminal fragment of polycystin-1 is mediated by a tethered peptide agonist"

SUPPLEMENTARY INFORMATION

**Table S1. Primer Sequences for PCR Cloning and Mutagenesis**.

| Primer | Sequence |
|---|---|
| 5’-CD5 Eco | 5’- TTCTAGAATTCCCTCGACCTCG -3’ |
| 3’-CD5-BsmBI | 5’- GACTAGCGTCTCATGCCTAGCACGGAAGC -3’ |
| mCleavStalkBsm For | 5’- GACTAGCGTCTCAGGCACTGCCTTCGGTGCC-3’ |
| mΔStalkBsmFor | 5'- GACTAGCGTCTCAGGCAGTGCAAGCATCAACTACATTGTCC -3' |
| TMI-EcoRV | 5’-GACTAGGATATCCCTCTGGACTCTAGTAAAGCG-3’ |
| hCTF Xho-BsmBI For | 5’-GACTAGCTCGAGCGTCTCAGGCACCGCCTTCGGCGCCAGCCTCTTC-3’ |
| BsrGIstalk-Rev | 5’- AGGGTCTGGGTAGAGTGCTT -3’ |
| hG3052R Top | 5’-CACCGCCTTCCGCGCCAGCCTC-3’ |
| hG3052R Bott | 5’-GAGGCTGGCGCGGAAGGCGGTG-3’ |
| hV3057M Top | CGG CGC CAG CCT CTT CAT GCC CCC AAG CCA TG |
| hV3057M Bott | CAT GGC TTG GGG GCA TGA AGA GGC TGG CGC CG |
| hR3063C For | 5’-GCCCCCAAGCCATGTCTGCTTTGTGTTTCCTGAG-3’ |
| hR3063C Rev | 5’-CTCAGGAAACACAAAGCAGACATGGCTTGGGGGC-3’ |
| hE3068D Top | GTC CGC TTT GTG TTT CCT GAT CCG ACA GCG GAT GTA AAC |
| hE3068D Bott | GTT TAC ATC CGC TGT CGG ATC AGG AAA CAC AAA GCG GAC |
| hP3069L Top | CCG CTT TGT GTT TCC TGA GCT GAC AGC GGA TGT AAA CTA C |
| hP3069L Bott | GTA GTT TAC ATC CGC TGT CAG CTC AGG AAA CAC AAA GCG G |
| hN3074K Top | GCC GAC AGC GGA TGT AAA GTA CAT CGT CAT GCT GAC |
| hN3074K Bott | GTC AGC ATG ACG ATG TAC TTT ACA TCC GCT GTC GGC |
| mG3044R Top | CTA GGC ACT GCC TTC CGT GCC AGC CTT TTT G |
| mG3044R Bott | CAA AAA GGC TGG CAC GGA AGG CAG TGC CTA G |
| mQ3055C Top | GTG CCT CCC AGC CAT GTC TGC TTC ATC TTT CCT GAA CCA TC |
| mQ3055C Bott | GAT GGT TCA GGA AAG ATG AAG CAG ACA TGG CTG GGA GGC AC |

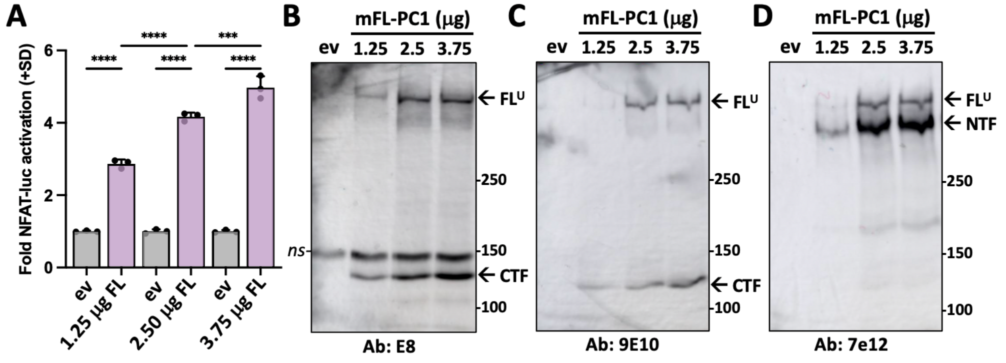


**Fig S1. Constitutive activation of NFAT-Fluc by FL-PC1 is 'dose-dependent'**. (**A**) Cells were transfected with 1.25. 2.5, or 3.75 ug of mFL-PC1 construct or the equimolar amount of empty expression vector (ev). Graph shows means +SD (n = 3) for activation of NFAT reporter relative to the ev control for each amount of FL-PC1. (**B,C**) Western blots of duplicate gels run with total cell lysates from transfection in (A) probed with E8 (B) or 9E10 (C). Lane ev from ev control for 3.75 ug FL-PC1. (**D**) Blot from (B) was stripped and reprobed with 7e12. $FL^U$, NTF, and CTF bands of PC1 are indicated. *ns*, nonspecific band.

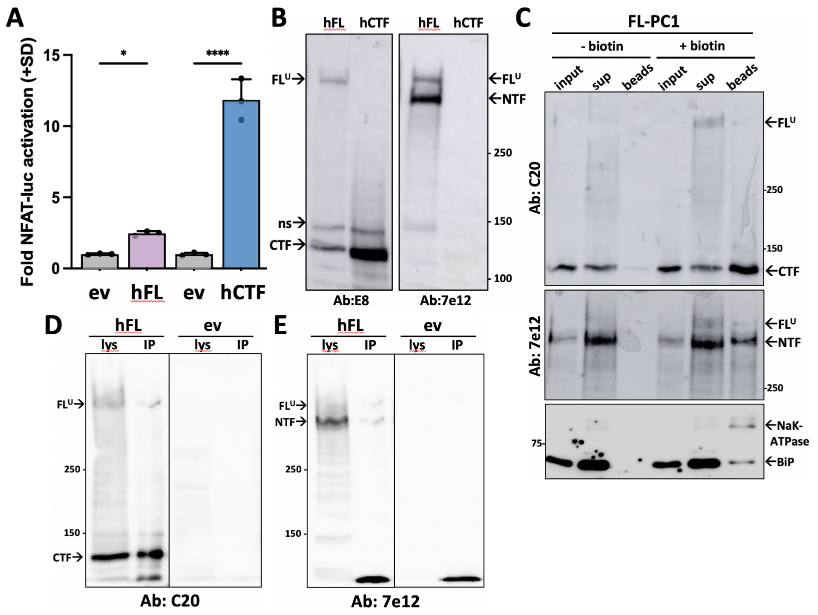


**Fig S2. Human (h) FL-PC1 and CTF also have differential abilities to activate the NFAT reporter.** (**A**) NFAT reporter activation by 2.5 ug of hFL-PC1 (hFL) and 75 ng of hCTF expression constructs expressed relative to each construct's equimolar empty vector (eV) negative control (gray bars). Shown are means + SD for 3 technical replicates. SD, standard deviation. Results are representative of 5 independent experiments. (**B**) Western blot of total cell lysates from the transfection in (A) probed with E8, then stripped and reprobed with 7e12. Normalization of NFAT reporter activation to the relative levels of total E8-reactive bands yields a 6-fold activation by CTF over corresponding ev. ns, non-specific. (**C**) Western blot of representative surface biotinylation assay with hFL-PC1-transfected cells. NHS-biotin was omitted from the procedure for half of the transfected cells (i.e., - biotin lanes). Aliquots of the starting cell lysate (input; before neutravidin pulldown), supernatant (sup; following neutravidin bead removal), and biotinylated proteins bound to neutravidin beads (beads) were electrophoresed on a 4.2% polyacrylamide-denaturing gel, blotted, and probed with C20 antibody. The blot was then stripped and reprobed with 7e12 antibody. Similarly, additional aliquots of input, sup and beads were run on a 7.5% polyacrylamide-denaturing gel, blotted and probed with anti-BiP and then reprobed with anti-Na,K-ATPase antibodies (Ab) as markers of ER and plasma membrane proteins, respectively. The – biotin beads lane demonstrates the specificity of the biotinylation analysis.

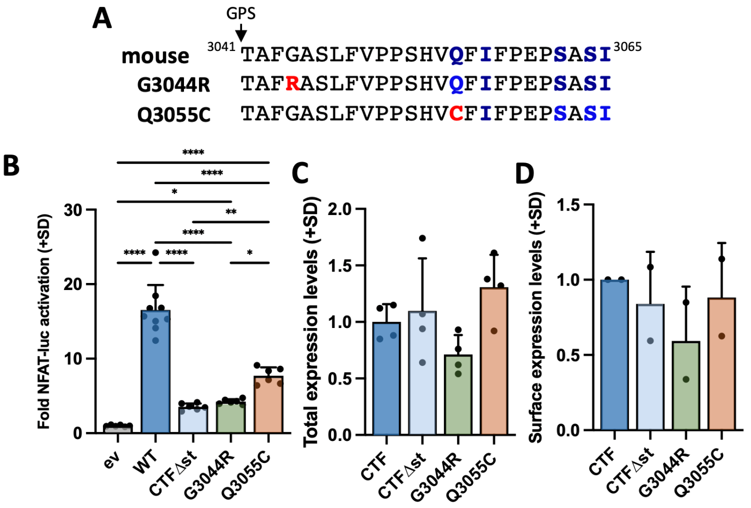


**Fig S3. ADPKD-associated missense mutations in mouse PC1 CTF effect signaling to NFAT reporter**. (**A**) Stalk region sequence of mouse PC1 CTF and two orthologous ADPKD mutations (G3044R, Q3055C; in red). Blue residues indicate those that differ between human and mouse. Missense mutation residues shown in bold red. (**B-D**) Cells were transfected with empty expression vector (eV), wild type (WT) CTF, or CTF mutants G3044R and Q3055C for NFAT reporter activation (B) or surface biotinylation assays (D). Western blots were performed with lysates from the signaling (total expression) and biotinylation beads (surface expression) and analyzed by densitometry to determine total and cell surface levels of the mutants relative to WT CTF protein (C,D). Graphs show means + SD, where n=3 replicates in each of 3 separate experiments, except for surface levels in (D) where n = 1 in each of 2 separate experiments. SD, standard deviation.
